# Statistics of Cellular Evolution in Leukemia: Allelic Variations in Patient Trajectories Based on Immune Repertoire Sequencing

**DOI:** 10.1101/037770

**Authors:** Hong Gao, Chunlin Wang, Junhee Seok, Marcus W. Feldman, Wenzhong Xiao

**Affiliations:** Stanford Genome Technology Center and Department of Biochemistry, Stanford University, Stanford, California, United States of America; Department of Biology, Stanford University, Stanford, California, United States of America; Department of Surgery, Shriners Burn Center and Massachusetts General Hospital, Harvard Medical School, Boston, Massachusetts, United States of America

**Author notes:** Current Address: Department of Bioinformatics, Illumina Inc., Santa Clara, California, United States of America (Hong Gao). Current Address: Sirona Genomics, Mountain View, California, United States of America (Chunlin Wang). Current Address: Department of Electrical Engineering, Korea University, Seoul, South Korea (Junhee Seok).

## Abstract

The evolution of a cancer system consisting of cancer clones and normal cells is a complex dynamic process with multiple interacting factors including clonal expansion, somatic mutation, and sequential selection. As a typical example, in patients with chronic lymphocytic leukemia (CLL), a monoclonal population of transformed B cells expands to dominate the B cell population in the peripheral blood and bone marrow. This expansion of transformed B cells suggests that they might evolve through processes distinct from those of normal B cells. Recent advances in next generation sequencing enable the high-throughput identification and tracking of individual B cell clones through sequencing of the V-D-J junction segments of the immunoglobulin heavy chain (IGH). Here we developed a statistical approach to modeling cellular evolution of the immune repertoire. Adapting the infinitely many alleles model from population genetics, we studied abnormalities occurring in the immune repertoire of patients as substantial deviations from the null model. The Ewens sampling test (EST) distinguished the immune repertoires of CLL patients with imminent relapse from healthy controls and patients in sustained remission. Extensive simulations based on sequencing data showed that EST is sensitive in detecting cancer-related derangements of the IGH repertoire. In addition, we suggest two potentially useful parameters: the rate at which donor’s B cell clones enter the circulation and the average time to regenerate a transplanted immune repertoire, both of which help to distinguish relapsing CLL patients from those in sustained remission and provide additional information about the dynamics of immune reconstitution in the latter patients. We anticipate that our models and statistics will be useful in diagnosis and prognosis of leukemia, and may be adapted for application to other diseases related to adaptive immunity.

## Introduction

Since the 1970s, cancer has been known to be a complex cellular evolutionary process [1-3]. In a cellular system with cancer (cancer system, for short), cancer cells evolve through processes distinct from normal cells. Analogously to the evolution of human populations, a cancer system may experience clonal expansion, somatic mutations, selection pressure, competition between cancer clones and normal cells, as well as bottleneck effects caused by treatment [1-3]. Evolution of cancer systems has recently become a focus of research, as it holds the potential for uncovering cellular disease mechanisms and for developing effective therapies for cancer patients. However, few studies to date have applied evolutionary theory to cancer biology for diagnosis and prognosis of cancer patients [2-5]. Here we apply evolutionary models and genetic statistics to understand the dynamics of cell development and evolution in a cancer system. This analysis may potentially aid in cancer diagnosis and prognosis.

In this study, we focus on chronic lymphocytic leukemia (CLL) as an example of a blood cancer system. CLL is a malignancy of mature B cells, where a monoclonal population of transformed B cells dominates the B cell population in a patient. It is the most common adult leukemia in the United States with approximately 15,500 new cases and 4,400 deaths per year [6]. Conventional immuno-chemotherapy may prolong survival, but is not curative. Allogeneic hematopoietic cell transplantation (allo-HCT) can provide long-term remission, and possible cure, for roughly half of patients with high-risk CLL [7-8]. Unfortunately, about 40 percent of CLL transplant patients ultimately relapse [9-11]. Quantification of minimal residual disease (MRD) after chemotherapy or transplantation tracks the frequency of cancer clones in a patient’s immune repertoire, and allows clinicians to monitor the efficacy of treatment by providing early detection of CLL progression at the molecular level [9-11].

The immune repertoire of an individual is the collection of antigen receptors of B or T cells in the peripheral blood at a given time [12-20]. In the B cell compartment, each antigen receptor gene is generated through rearrangement of the variable (V), diversity (D), and joining (J) gene segments of the immunoglobulin heavy chain (IGH) locus and V-J rearrangement of the kappa or lambda light chain loci. Due to recombination of V, D and J gene segments and random nucleotide changes at the junction of those gene segments, the number of potential antigen receptors is enormous. The immune repertoire of healthy subjects is generally quite diverse, while loss of immune repertoire diversity has been found to be associated with aging and various diseases including cancer [21-24].

Next generation sequencing (NGS) provides a cost-effective opportunity to sequence the V-D-J gene segments of individual B cells [5,18,25-27]. As each V-D-J sequence is a unique identifier for each B cell, NGS allows us to monitor B cell evolution in an immune repertoire and enables investigation of methods for detecting rapid clonal expansion of malignant B cells at the expense of the normally diverse B cell repertoire. This approach may ultimately improve cancer diagnosis and prognosis; however, the lack of sophisticated statistical and computational approaches for interpreting such NGS data currently limits its use in clinical decision-making.

In this study, we develop a statistical approach to modeling the cellular evolution of the immune repertoire. This approach adapts the infinitely many alleles model from classical population genetics theory [28-30]. The null model describes the variability of a healthy immune repertoire in the absence of antigen stimulation, and deviations from this null model may reveal disequilibrium or abnormality in an immune repertoire, indicating potential risk of disease. The Ewens sampling test (EST) [28,29,31] can be applied to identify deviations from the null model. We performed simulations to demonstrate that EST can detect irregularity in the immune repertoire with relatively high sensitivity. We then applied EST to our IGH V-D-J sequencing data from CLL patients [27] and showed that EST distinguished the immune repertoires of CLL patients with active or imminent relapse from those of healthy individuals and patients in long-term remissions (i.e., lasting well beyond the period covered by our sequencing study). Additionally, we estimated two parameters of interest for patients undergoing allo-HCT: the rate at which donor novel B cells enter the circulation and the average time to reconstruct the transplanted immune repertoire after transplant. These characterize the evolutionary trajectories of the immune repertoire following allo-HCT. As the approach is able to detect deviations of an immune repertoire from normal, we anticipate that it can be applied to other types of B cell or T cell cancers as well as other diseases related to adaptive immunity.

## Results

In order to distinguish the cellular evolutionary pattern of patients with CLL from controls, it is essential to first model the evolution of healthy immune repertoires and then to use this as a null model to develop statistical tests. In examining the frequency spectra of V and J gene rearrangements from CLL patient and donor samples [27], we observed high diversity in healthy donors (Fig 1(A)) and a highly skewed frequency distribution, usually with one dominant clone, in CLL patients (Fig 1(B)). We modeled evolution of the immune repertoire with the Moran infinitely many alleles model assuming a constant population size (see below for justification), and applied the Ewens sampling test to detect CLL as a significant deviation from the normal pattern. We also estimated the incorporation rate of novel B cells into the immune repertoire as an indicator of disease status. Next we modeled the development of the transplanted immune repertoire (the fraction of the recipient’s B cell repertoire derived from the transplanted donor cells), whose size varies over time, using a linear birth process with immigration to relax the assumption of constant population size in the infinitely many alleles model. Based on this process, we estimated the average time to generate the observed transplanted immune repertoire and used it as a measure of expansion or shrinkage of the transplanted repertoire.

**Fig 1.**
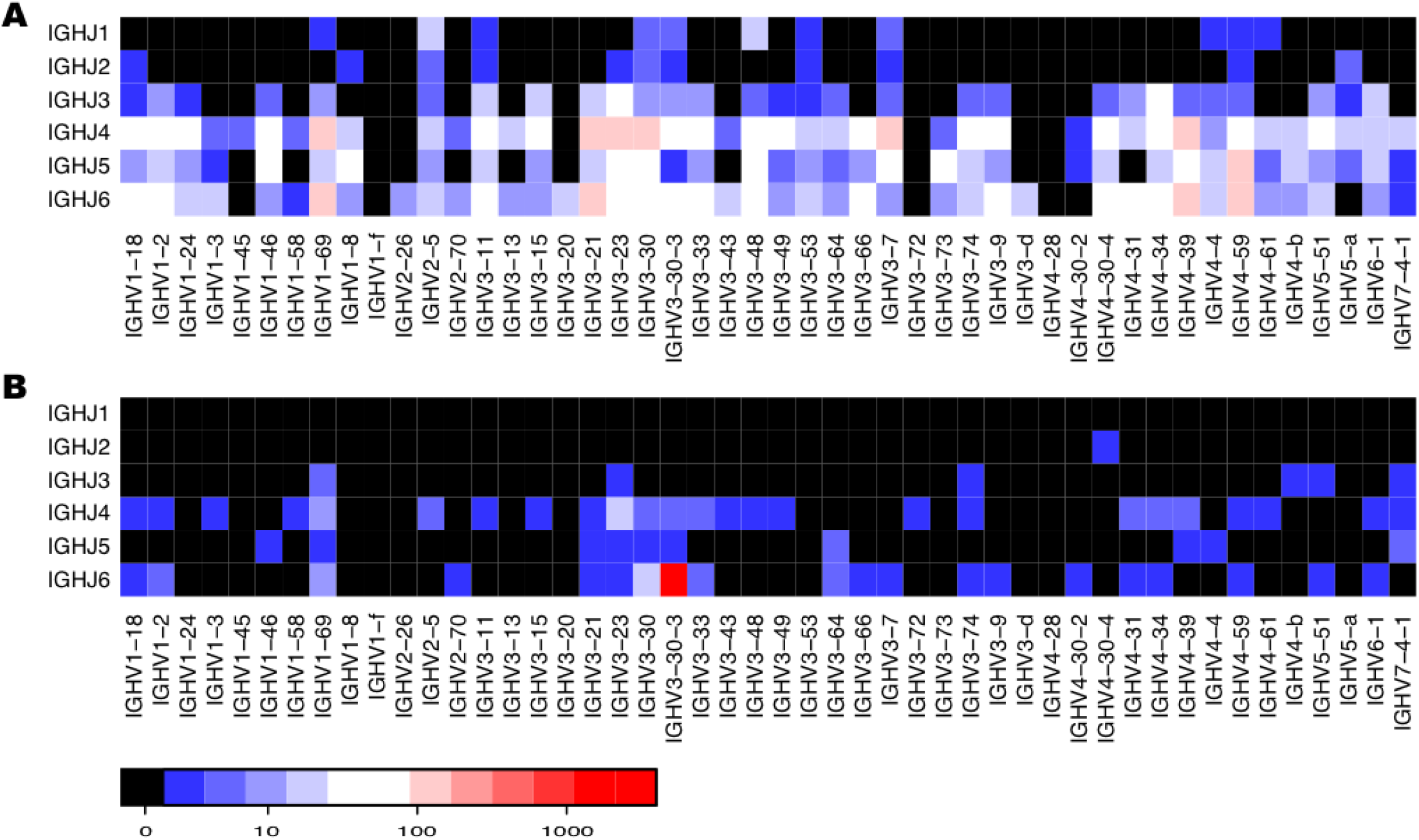
Heatmap of frequency spectra of V-J combinations in immune repertoires. Frequency spectra of V-J combinations in immune repertoires of a healthy donor (A) and a CLL patient SPN4077 at diagnosis (B) across 48 functional V segments (x axis) and six functional J segments (y axis). Color indicates the number of cells on the log scale.

### I. Modeling cellular evolution in the immune repertoire

Motivated by the distinct diversity patterns of CLL and donor samples, and to understand MRD and immune reconstitution after allo-HCT, we adapted an evolutionary stochastic process, the infinitely many alleles model [28-30], to model the generation and elimination of B cells. The Moran version of this model describes a neutrally evolving population in which alleles are created or lost through a birth and death process. To apply this model, we made a simplifying assumption that after novel B cells acquire VDJ rearranged antigen receptors and mature in bone marrow or lymph nodes, they enter the circulation and proliferate independently and spontaneously in the absence of antigen stimulation, as illustrated in Fig 2 (A). Some of the original B cells in the population may be eliminated, some may remain unchanged, and others may proliferate. Since mature B cells’ half-life is approximately 5-6 weeks [32], when looking backward, the immune repertoire pattern at any given instant is assumed to have been generated from B cells entering the circulation within the prior three months.

**Fig 2.**
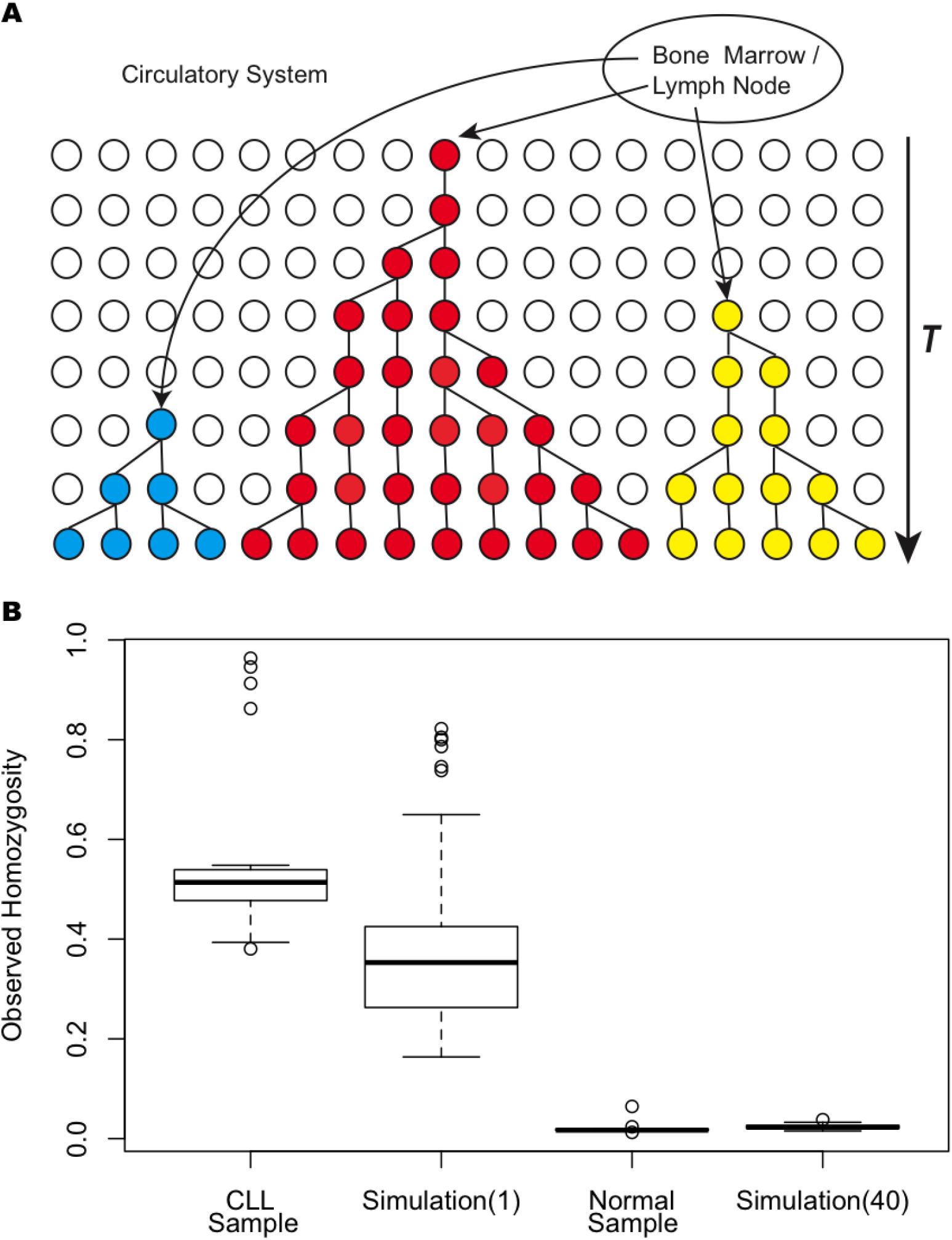
Illustration of B cell evolutionary model (A) and evaluation of model fitness by forward simulation (B). (A) Schematic illustration of the general developmental process of B cells over time *T.* After maturing in bone marrow or training in lymph nodes, B cells represented by circles enter the circulation to proliferate and initiate clones independently and spontaneously. Each color represents a distinct clone and the distinct clones enter the circulation at different times. (B) Distributions of observed homozygosity of data from patient/normal samples and forward simulations. From left to right, respectively, are represented CLL samples, forward simulation with incorporation rate *θ*= 1, normal (healthy) donor samples, and forward simulation with *θ*= 40.

To simplify the modeling of B cell development in allo-HCT patients, we made several additional assumptions. First, we assumed a constant population size of B cells in an immune repertoire, since the number of B cells is generally stable in the human circulation (a normal B-cell count is between 50 and 500 per microliter [33]). Second, the process of generating a B cell includes multiple transitional states, progenitor B cell, pro-B cell, pre-B cell, transitional B cell, naive B cell, plasma blast, and memory B cell. Also the majority of B cells in the blood are naïve B cells [34]. Here we reduce this process to two states according to their location, inside and outside the circulation. Third, in our context, a novel B cell represents a normal B cell with a distinct IGH V-J combination since it is difficult to reliably determine the D gene segment due to its short length in some cases (it is straightforward to generalize this model to V-D-J combinations or even clonotypes if these can be accurately identified). Fourth, we assumed that a new V-J combination is randomly generated from 288 combinations of 48 functional V and six J gene segments. Although the total number of possible V-J combinations, 288, is finite, we assumed that it is sufficiently large in a population of 15,000 B cells to approximate the generation of previously unseen mutant alleles (new clones), as assumed in the infinitely many alleles model [28-30]. Last, the unit of time for this system is assumed to be the average time interval between two consecutive proliferation events of a B cell.

The evolution of B cells in the immune repertoire in the absence of antigen stimulation is very similar to the genetic drift of alleles in a constant-size neutral population. Novel B cells enter the circulation following maturation outside the circulation (i.e., in the bone marrow or lymph nodes). This is analogous to the process of introducing new allelic mutants into the population. Due to allelic exclusion, each B cell carries only one functional copy of the IGH gene [35], so for *M* cells there are *M* copies of the gene. We then model the evolution of the immune repertoire with the Moran infinitely many alleles model, where alleles are generated or lost through a birth and death process [29,36,37]. Theoretically, there exists a stationary distribution of allelic configurations for this model, which is the Ewens sampling distribution (ESD) [29,36]. The rate at which novel B cells enter the circulation is equivalent to the mutation rate of alleles, denoted by *μ.* We let *θ*=*2Nμ*, where *N* is the constant B cell population size in an immune repertoire. *θ*/*2* can be interpreted as the expected number of novel B cells entering the circulation per unit of time. Henceforth, we refer to *θ* as the incorporation rate of novel B cells in the circulation.

To examine whether this model reflects evolution of the immune repertoire, we performed forward simulations to evaluate whether the Ewens distribution fits the clonal frequencies of an empirically determined immune repertoire. We simulated B cell clones following the modified Polya urn model [38] with different values of the incorporation rate *θ*, as ESD arises naturally from the modified Polya urn model (see Methods section III). When *θ* is large, e.g. 40 (S1 Fig (B)), which means 20 novel B cells are expected to enter the circulation each unit of time, the frequency spectrum of V-J combinations resembles that of the normal immune repertoire with relatively high diversity (Figs 1 (A) and 2 (B) normal). As *θ* decreases, the diversity decreases substantially. When *θ* is 1 (S1 Fig (A)), the frequency pattern is very close to that of the CLL repertoire at diagnosis with a low diversity (Fig 1 (B)). We repeated the simulation 100 times for each value of *θ* and estimated the observed homozygosity for each simulation. Fig 2 (B) shows that the distribution of observed homozygosity for simulations with *θ*=1 overlaps that of cancer (CLL) samples and the distribution of homozygosity simulated with *θ*=40 covers that of donor samples. This simulation shows the consistency of our model with the biology of lymphocytes. When the incorporation rate *θ* of novel B cells is low, a higher fraction of identical cancer B cells are generated in the system. When *θ* is high, the majority of B cells entering the circulation are novel, leading to high diversity in the immune repertoire. The simulation suggests that our model may be a reasonable description of B cell evolution in the immune repertoire.

### II. Application of Ewens sampling test

The stationary allelic configuration distribution of this infinitely many alleles model in the absence of any selective process, namely the Ewens sampling distribution [29,36], can describe the V-J combination frequency spectrum in a snapshot of a healthy immune repertoire. When cancer clones rapidly expand to dominate the repertoire, the diversity is substantially reduced and the pattern of variation in the repertoire diverges from that expected from the ESD. This led us to apply the Ewens sampling test [29,36] to detect whether the V-J combination frequency spectra of CLL patient immune repertoires differ from normal.

In a previous study [27], we sequenced IGH variable regions in serial samples of six CLL patients up to two years (740 days) after transplantation (Stanford Patient Numbers SPN4077, SPN3860, SPN3751, SPN3873, SPN3740, and SPN3975), and six corresponding donor samples (see Methods section V). Of these patients, SPN3740 and SPN3975 experienced early CLL relapse at 182 and 210 days following allo-HCT, respectively. Three patients remained in long-term remission until day 1,120 (SPN3751), day 980 (SPN 4077), and day 798 (SPN3873). SPN 3860 had not relapsed with 1,810 days of follow-up. SPN3740, SPN 3751, SPN3873, and SPN4077 received Rituximab at 56, 63, 72, and 79 days post transplantation, which eliminates the vast majority of mature B cells in circulation with an average duration of response of six months. Thus, samples collected at 180 days post HCT are excluded from the analysis, due to the potential confounding effects of Rituximab treatments. We acquired roughly 15,000 functional reads from each sample after eliminating non-functional alleles (i.e., those with premature stop codons) from our analysis. Henceforth, we assume each functional read corresponds to one B cell.

After identifying germline V and J gene segments corresponding to each read, we examined the V, J gene usage pattern for each sample. The temporal V-J frequency spectra of patient SPN4077 were shown in Figure 5 of [27] and here in Fig 1. At diagnosis (Fig 1 (B)), the highly skewed frequency distribution with one dominant clone suggests heavily reduced diversity of the underlying immune repertoire. Over a period of one year after allo-HCT, the proportion of the dominant cancer clones gradually diminished while the nascent clones expanded steadily. After one and a half years, the pattern of the recipient’s repertoire was as diverse as that of the donor (donor’s repertoire pattern is shown in Fig 1 (A)). Supporting Figure S5 in [27] shows the temporal trajectories of the spectra of V-J combinatorial diversity for the other five patients. Among them, the spectra of patients SPN3975 and SPN3740 with early relapsed CLL remain highly skewed over time.

We then applied EST to the immune repertoire sequencing data of these patient samples. As illustrated in Fig 3 (A,B), all the donors’ immune repertoires follow the null distribution with p-values of EST approximately 1, and their observed homozygosities (the test statistic of EST) are close to 0, indicating a high diversity (Fig 3 (C,D)). However, the immune repertoire of CLL patients at diagnosis (SPN4077) or after relapse (SPN3740 and SPN3975) reject the null hypothesis (Fig 3 (B)), as they generally have a highly skewed frequency distribution of B cell clones. When the patients recover after allo-HCT, their repertoire patterns follow ESD again. For patient SPN4077 in Fig 3 (A), the p-value at 56 days post allo-HCT is the same as that at diagnosis, as the cancer clone remained dominant although the transplanted B cells began to expand. At one year, novel B cell clones dominated SPN4077’s immune repertoire with the observed homozygosity close to 0 and the p-value reaching 1. The observed homozygosity of immune repertoire of patients SPN3860 and SPN3751 reaches a level similar to healthy donors at 18 months post allo-HCT (Fig 3 (C)) and their p-values are close to 1 (Fig 3 (A)). The repertoire homozygosity of patient SPN3873 remains higher than donors’ levels, and the p-values are significant for the three time points after HCT, implying this patient may be on the verge of relapse. This result suggests that this test, based on immune repertoire diversity, can reflect the MRD status of patients, and has the potential for prognosis of the allo-HCT treatment for these patients.

**Fig 3.**
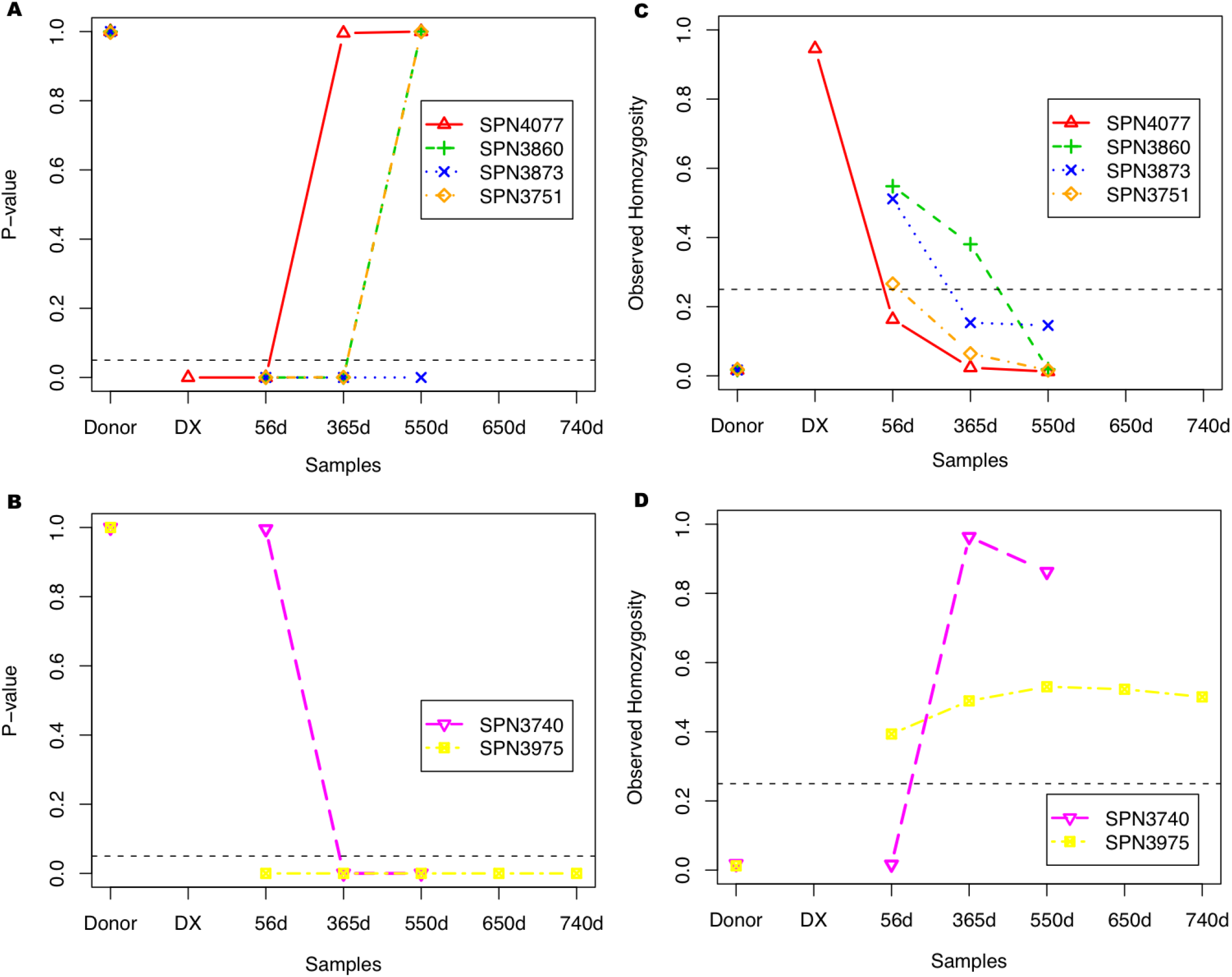
Temporal variation of p-values (A-B) and test statistic (C-D) of EST for deviations from the normal pattern in Fig 1 (A) among samples of six patients. SPN4077, SPN3860, SPN3873 and SPN3751 were in sustained remission following allo-HCT (A and C); while SPN3975 and SPN3740 had imminent relapsed CLL post-HCT with p-values close to zero (B and D). All the donors have p-values 1. The black dashed line in A and B represents the significance level of 0.05 for EST. The black dashed line in C and D is an arbitrary separation of patients for observed homozygosity. Note: for SPN4077 data are available at four time points, while only three time points are available for SPN3860, SPN3873 and SPN3751. The x-axis labels “Donor” and “DX” represent donor sample and patient sample at diagnosis, respectively. “56d”, “365d”, “550d”, “650d” and “740d” represent patient samples collected at different time points (in days) after allo-HCT.

We performed simulations to evaluate the sensitivity of our test in this application. We randomly simulated data-sets mimicking experiments that spike cancer clones into donor samples with the surrogate cancer clone proportions ranging from 1% to 99% of the whole repertoire (see Methods section IV). As illustrated in Fig 4 (A), the p-values for EST are close to 1 when the percentages of cancer clones are small. Then the p-values drop sharply as the percentages of cancer clones increase to approximately 15%. Though EST is known to be conservative it can accurately distinguish normal from abnormal patterns in the immune repertoire. The value of *θ* can be estimated from the frequency configurations as shown in Methods section I. The maximum likelihood estimate (MLE) of *θ* is denoted by 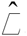.

**Fig 4.**
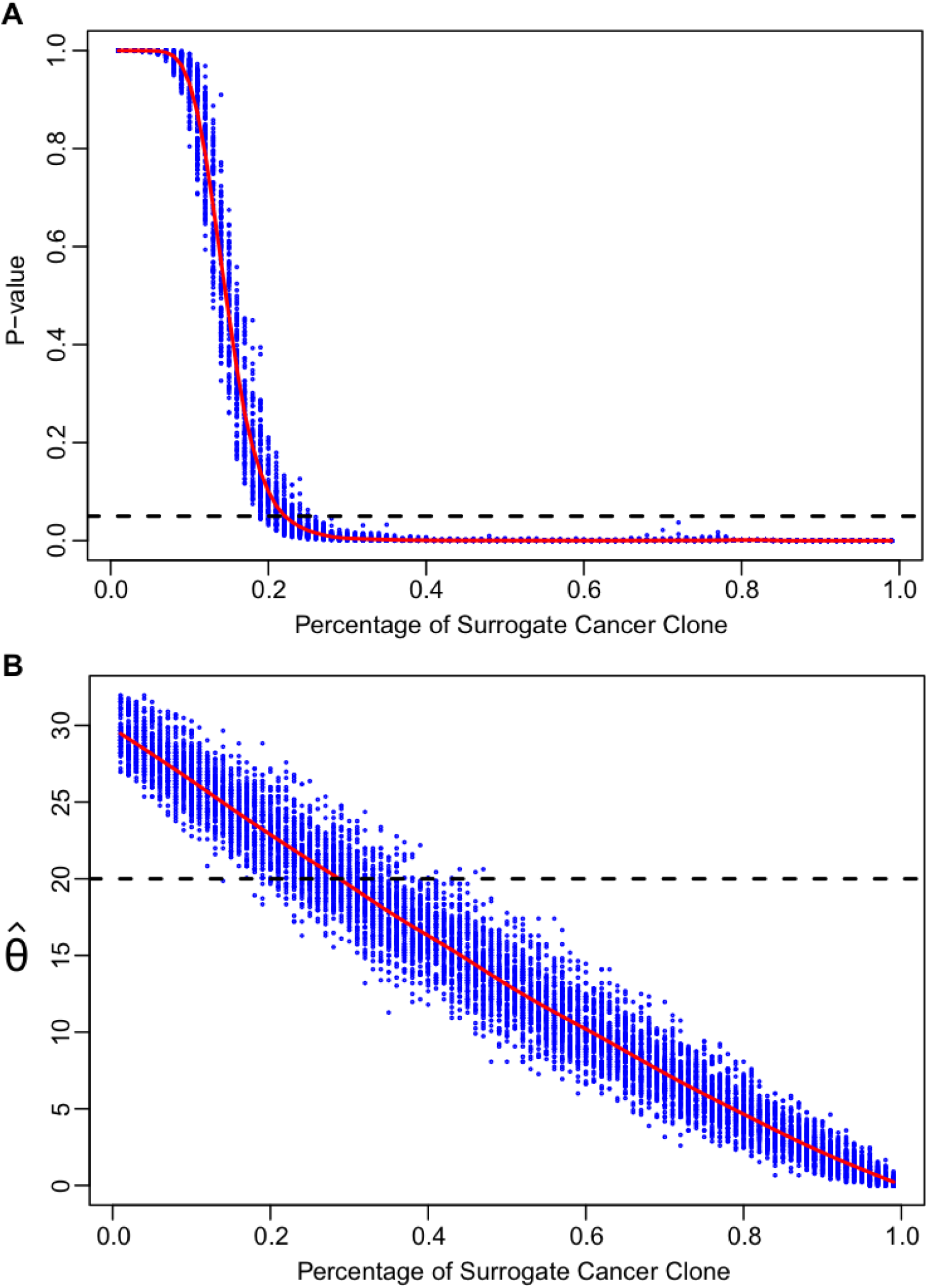
Sensitivity analysis of statistics based on the simulated data. The sensitivity of Ewens sampling test (A) and 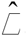 (B), MLE estimator of incorporation rate *θ*, respectively, at various abundances of surrogate cancer clones. The x-axis represents the percentage of the surrogate cancer clones in the entire immune repertoire. The black dashed lines represent the significance level of 0.05 for EST (Fig 4 (A)) and the *ad-hoc* cut-off 20 for 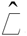 (Fig 4 (B)), respectively.

### III. Estimation of incorporation rate of B cells in immune repertoire

Addition of novel B cells can be viewed as a process of novel B cells migrating into the circulation. These novel B cells have an allele frequency distribution characterized by an incorporation parameter *θ*, which controls the dynamical status of the immune repertoire. We assume that in a healthy repertoire, *θ*/*2* novel B cells are expected to enter the circulation per unit time of the system. When cancer clones dominate the repertoire, very few novel B cells appear in the circulation. Thus incorporation events may be limited and *θ* becomes relatively small suggesting that 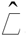 can be used to monitor the status of the immune repertoire. *θ* represents the rate at which novel B cells enter the system and the number of different B cell types (different V-J combinations in our simplified scenario) in a given immune repertoire is a sufficient statistic for *θ* [29,36,37].

We estimated *θ* from the simulated data sets and for the six patients using 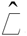 as described in Methods. The variation of 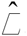 with the abundance of surrogate cancer clones is shown in Figure 4B. We chose an *ad-hoc* cut-off of 20 for 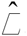, which clearly separates the trajectories of patients with imminent relapse from those with sustained remission. In patients SPN4077, SPN3860, SPN3751, and SPN3873 (Fig 5 (A)), we observed an overall increasing temporal trend of 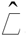 until it reaches the control level, implying rapid incorporation of B cells into the circulation. In contrast, this estimator is substantially lower in patients SPN3740 and SPN3975 (Fig 5 (B)), who had CLL relapse during the period, probably due to ineffectiveness of allo-HCT or Rituximab and other anti-B cell therapies in these patients. These results suggest that 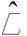 has the potential to distinguish normal samples from the cancer-like ones, and can serve as information supplementary to EST.

**Fig 5.**
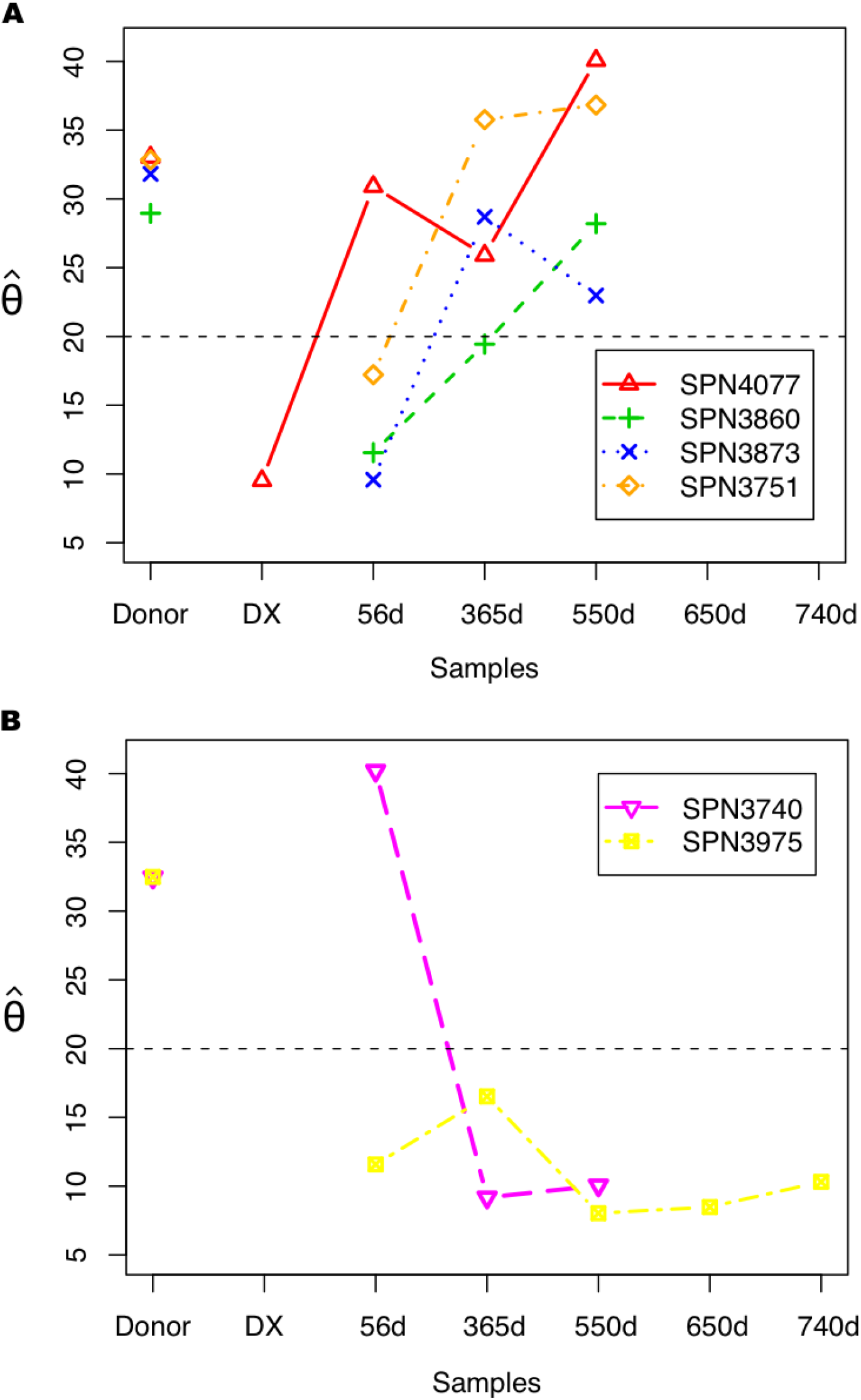
Temporal changes in estimated immigration rates 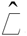 of novel B cells among patient samples. The grouping of patients and the x-axis labels are the same as in Fig 3. The black dashed line represents the *ad-hoc* cut-off of 20 for the estimated incorporation rate 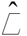.

### IV. Estimation of the average time to reconstitute immune repertoire

We also estimated the average time to generate the transplanted immune repertoire observed at a given time after removing any cancer clones and subclones. For the transplanted immune repertoire, we applied a different model, the linear birth process with immigration [29,36,37]. The reason for using this model is that the infinitely many alleles model assumes a constant population size, which may not apply if the size of the transplanted immune repertoire changes over the time. This linear birth process with immigration relaxes the assumption of constant population size, and may be appropriate to model the evolution of the transplanted immune repertoire.

In this model, we assume that the donor’s B cells with new V-J combinations migrate into the circulation according to a continuous time, pure-birth Markov chain with the immigration parameter denoted as 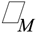. Upon migration, each B cell immigrant initiates a clone, the size of which follows a linear birth process with rate 1. Different clones evolve independently. Therefore, the population size of all the donor’s B cell clones is mathematically formulated as a linear birth process with immigration. Interestingly, the distribution of allelic configurations in the stochastic process at any given time is also the Ewens sampling distribution [29,36,37], as in the infinitely many alleles model. In the case of successful immune repertoire reconstitution, the donor’s B cell population, which starts very small, will grow following this process to a reasonable size—e.g. the normal number of B cells in the circulation—after which the population size will become stable with minor random fluctuations. In the case of cancer relapse, the donor’s B cell population either remains small or shrinks.

Using this model, we can estimate the expected time to generate a transplanted immune repertoire with a given size as shown in Methods section II. This is equivalent to estimating the expected age of the oldest donor-derived B cell clone in the repertoire, and is the sum of the expected time intervals between any two consecutive events, either immigration or proliferation events. This time parameter is of clinical interest, as it measures the expansion or shrinkage rates of the transplanted immune repertoire in the recipient’s circulation and can also be used to evaluate the patient’s recovery status after treatments. For our patient data, we estimated this time parameter for each sample from the six patients after treatment. Fig 6 shows that this time estimator increases monotonically for patients SPN4077, SPN3860, and SPN3873, who remain in remission for more than two years following allo-HCT, and decreases substantially for patients SPN3740 and SPN3975 with relapsed CLL during the same period, though Rituximab and other treatments likely also contribute to the sharp reduction. An exception is seen in patient SPN3751 for whom the estimated time of reconstitution drops sharply at one year, probably due to the prolonged effect of Rituximab, which eliminates most B cells over the period of six to nine months after allo-HCT; however, this drop is followed by remarkable increases. Generally, the estimated time to generate a transplanted repertoire reflects the recovery trend of patients and can be informative at early time points.

**Fig 6.**
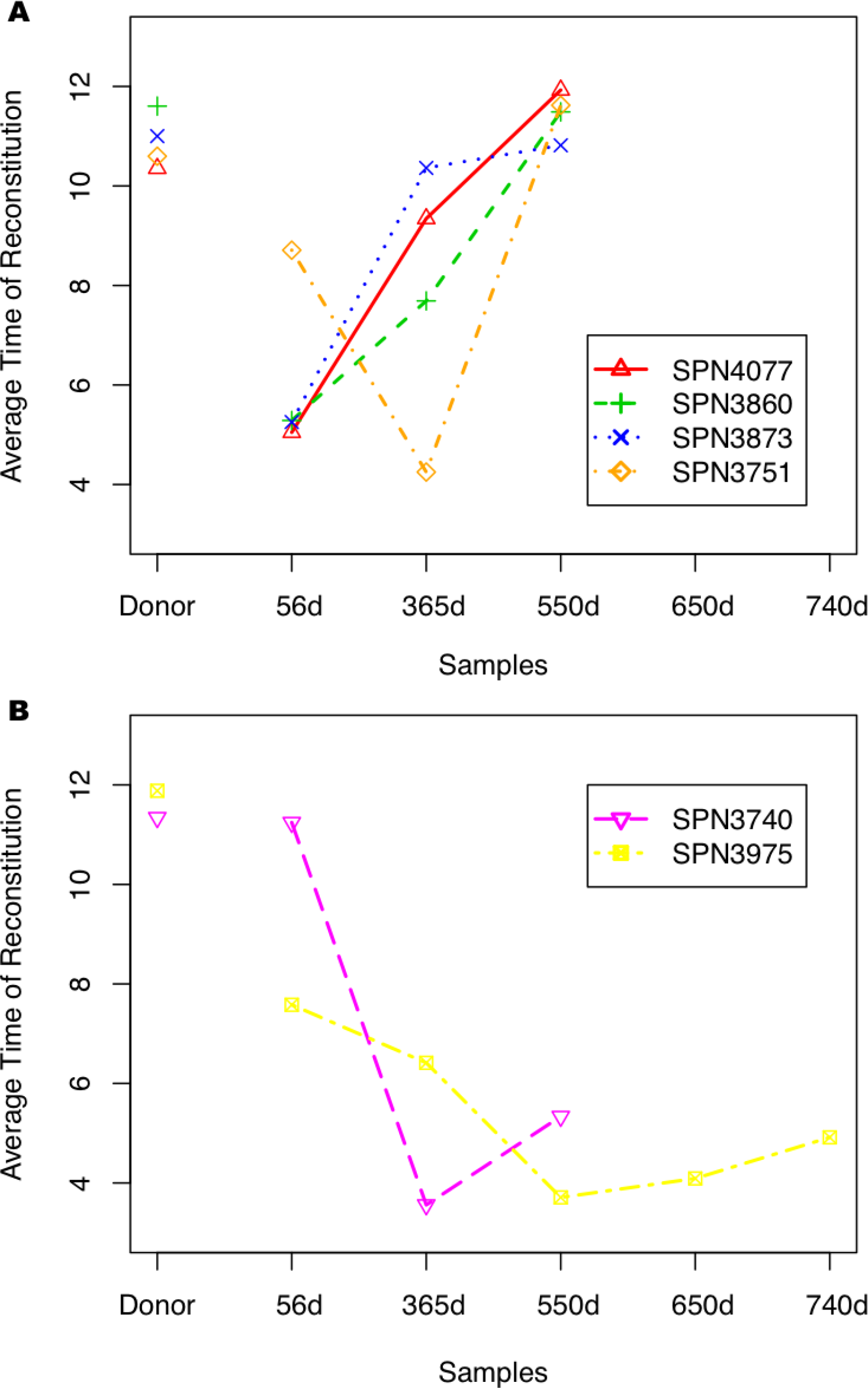
Temporal trend of estimated average time to construct transplanted immune repertoires in six patients. The grouping of patients and the x-axis labels are the same as in Fig 3.

## Discussion

There have been few studies to date that apply molecular evolutionary theories to understand the mechanisms underlying cancer systems, although cancer development is indeed an evolutionary process [1-4,39,40]. Maley et al. [4] used the Shannon diversity index to predict progression to esophageal adenocarcinoma, which inspired our current study. Here we applied the infinitely many alleles model to describe the spontaneous evolution of an immune repertoire in the absence of external selection. Any immune repertoire patterns that deviate significantly from the equilibrium of allelic configurations predicted by this neutral evolutionary model may be a result of the progression of blood cancers such as CLL.

Using IGH high-throughput sequencing data of CLL patient samples, we employed this model to investigate the reconstitution of the immune repertoire after allo-HCT. We applied EST to distinguish CLL patients from healthy individuals. We also estimated two parameters of clinical interest, the immigration rate of novel B cells into the circulation and the average time to generate the transplanted immune repertoire after allo-HCT. Applications to both experimental and simulated data demonstrated that our model mimics the real data well: statistics and tests derived from the model can successfully distinguish cancer samples from normal ones, and describe the evolutionary dynamics of an immune repertoire. In particular, these statistics are sensitive in distinguishing samples with immediate CLL relapses from those in remission for more than two years. They also revealed a potential negative effect of Rituximab post HCT, which functions as a bottleneck for a B cell population. Since Rituximab eliminates the majority of B cells, when administrating it to post-HCT patients, it necessarily leads to delays in the reconstitution of the normal immune repertoire.

Our statistics provide three additional insights. The EST assesses the diversity of an immune repertoire; the estimator 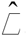 of incorporation rate reflects the evolutionary variation of a repertoire; and the estimated average time to repertoire reconstitution reveals the expansion or shrinkage of a transplanted repertoire. Together these metrics assess the quantitative properties of the immune repertoire in disease monitoring, including its diversity and dynamics. Thus, integrating these statistics may provide the potential for prognosis of patients with treatments. Large-scale studies are needed to confirm the applicability and accuracy of these statistical tools in patient care.

The application of our model and methods is not limited to blood cancer. By using analogous interpretations, it is straightforward to extend this approach to immune monitoring in other diseases, such as autoimmune diseases, infections and non-hematologic malignancies. The polymorphic variation in a healthy immune repertoire with mild fluctuations may often be described in terms of ESD. When diseases occur, the human body often generates strong and persistent immune responses against pathogens or other antigens and the immune repertoire generally exhibits highly skewed frequency spectra of lymphocyte clones, such as have been seen in colon cancer and systemic lupus erythematosus (unpublished data from Dr. C. Wang). Therefore, the diversity patterns of immune repertoires in these diseases may deviate from ESD and become detectable by our statistics.

Our stochastic model is a substantial simplification of the evolutionary mechanism of the actual dynamics of an immune repertoire and neglects many essential elements including positive and negative selection. It is predominantly applicable to rather ideal situations, such as in the absence of antigen stimulation or with mild or weak antigen stimulation, where the background (null) distribution follows ESD. Analysis of additional clinical samples will be needed to determine whether different repertoire signatures occurring in the context of clinical events such as infections will remain assessable with this method. To effectively model the response of the immune repertoire to other immunological diseases, the current model might be extended by incorporating parameters representing the selection pressures on specific lymphocyte clones. This would involve allowing lymphocyte clones, after entering the circulation, to proliferate at different rates, proportional to the strengths of antigen stimulation. By explicitly modeling positive and negative selection, the power to distinguish persistent abnormality in the immune repertoire from temporary fluctuations may be improved.

Currently our diversity modeling and analysis are mainly performed at gene segment levels, e.g., V and J gene segments, but these methods can be extended to the clonotype level. However, identification of different types of clonal sequences requires accurate identification of somatic hypermutations and sequencing errors. Lately extensive efforts have been made to improve sequencing techniques and reduce sequencing error rates. Currently Illumina sequencers produce approximately one sequencing error every 1000 nucleotides, while iontorrent machines produces ˜1 error every 100 nucleotides. Therefore, using Ilumina sequencers it is possible to estimate diversity metrics at clonotype levels, but they tend to produce multiple replicates for each clonotype. Thus, before applying our models, replicates need to be removed to comply with our assumption that each read corresponds to one cell. Advances in sample processing enable PCR-free sequencing, which eliminates the amplification step and will not produce replicates. Applying these new technologies will enable estimation of diversity metrics at clonotype levels, and permit inference concerning more subtle variations in patient immune repertoires.

## Methods

### I. Moran infinitely many alleles model and derived statistics

We model the evolution of the immune repertoire as a Moran infinitely many alleles model [41,42], which assumes that individuals in a population are created or lost through a birth and death process with a constant population size. In our case, B cells may proliferate or die randomly in a population. According to this model, the allelic configuration of the V-J combinations at a stationary state follows ESD and we can formulate a statistical test based on this null distribution. We choose the observed homozygosity as the test statistic and the p-value can be theoretically computed using Ewens sampling formula [41,42]. However, in our scenario, the sample size is generally large, on the order of 10^4^, thus it is impossible to enumerate all possible configurations of V-J combinations. Monte Carlo simulation is used to generate the p-values of EST, following Slatkin [41,42], and the significance cut-off is set at 0.05. The code for our application of the EST is available upon request.

To estimate the incorporation parameter *θ*, we directly apply Ewens’ result that the number of different allele types in the system, *K*, is a sufficient statistic for *θ*, and the MLE 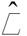 of *θ* is the solution of the equation 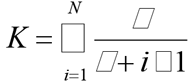 [29,36], where *N* is the population size and *0*<*K*< *N*. Then we use the bisection method to solve the above equation to obtain 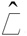 [41,42].

### II. Linear birth process with immigration and derived estimator

We show the derivation of linear birth process with immigration for the development of a transplanted immune repertoire. The number of novel B cells that have entered the circulation and initiated clones by time *t* follows the stochastic pure birth process *P*(*t*), which has a rate 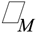 and initial value zero. The *i*-th novel B cell, which comes from the donor, migrates into the circulation at the time *T_i_* (*0* ≤ *T_1_* < *T_2_* <….) and then proliferates to initiate a clone independently from other clones according to a linear birth process with rate 1. We let 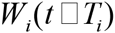 measure the size of the *i*-th B cell clone at time *t* after entering the circulation at the time 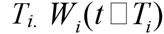 starts at *W_i_* (0) = 1 and increases at an infinitesimal birth rate *n_i_* (*t*) for 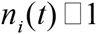, where *n_i_*(*t*) represents the size of the *i*-th B cell clone at time *t*. Thus the total size of the transplanted B cell population in an immune repertoire at time *t* is the sum of sizes of all the present B cell clones 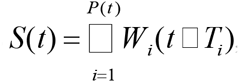, starting at *S(0)* = 0, with infinitesimal rates 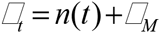 for 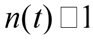, where 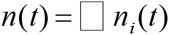 is the total size of all the B cell clones at time *t*. This process *S*(*t*)is called a linear birth process with immigration.

We then estimate the average time to generate the transplanted immune repertoire observed at a given time *t* by first identifying and removing cancer clones and then summing up the expected time intervals between any two consecutive events, either immigration or proliferation. For each sample of the six patients, we estimate this time parameter *T_M_* 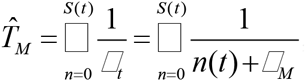, where *S(t)* is the transplanted B cell population size of that sample at time *t* and 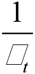 is the expected time until the next event takes place, which is the reciprocal of rate 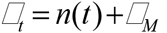 for 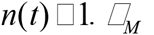 is estimated by solving 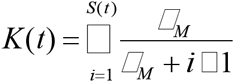, where *K*(*t*) is the number of different B cell types in that sample at time *t* [29,36,37].

### III. Forward simulation

To examine whether our model fits the immune repertoire data, we performed forward simulation following the modified Polya urn model (also known as the Chinese restaurant process) [38] with different values of the incorporation rate *θ*. At time *t* =1, the immune repertoire is empty and a novel B cell enters the system. At time *t* = *N*(*t*), the *N*(*t*)-*th* B cell is added to the repertoire either by an incorporation event with probability 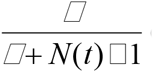 or by a proliferation event from the *i*-th existing B cell clone with probability 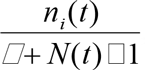, where *n_i_*(*t*) is the size of the *i*-th existing B cell clone at time *t* and 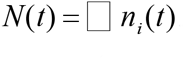. The simulation is stopped when the population size reaches 15,000. We used two values for parameter 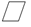, 40 and 1, corresponding to the estimated incorporation rate 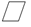 from donor samples and cancer samples, respectively. The simulation was repeated 100 times for each value of 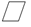 and observed homozygosity computed for each simulation and each patient sample.

### IV. Simulation for performance evaluation of test statistics

To evaluate the sensitivity of our methods, we performed extensive simulations mimicking the spiking-in of cancer clones at different proportions into control (donor) samples. We tested 99 levels of cancer clone abundance, ranging from 1% to 99% of the total read size. For each of the 99 proportions, we randomly chose one of the control samples and then picked one clone as the surrogate cancer clone. Next, we assigned a new clone size calculated according to the proportion assigned to this cancer clone and sampled the remaining clones without replacement to constitute the residual immune repertoire, scaling the total sample size to 15,000. We repeated this procedure 100 times per proportion level, that is, 9,900 simulations in total.

### V. Immune repertoire sequence data from CLL patients

We have previously collected serial blood samples from six CLL patients at diagnosis and after allo-HCT at days +56, +365, and +550. (Samples collected at day +180 used in [27] were removed as those patients were treated with Rituximab at three months post-transplant, which may confound analyses.) Patient SPN3975 has two samples collected at additional time points at days +650 and +740. The samples of six donors were also collected to serve as references. These samples were then sequenced by Roche/454 technology. After initial filtering, we mapped each read to germline sequences from ImMunoGeneTics (IMGT) database [43] using the Asymmetric Smith-Waterman algorithm [44] to identify the germline types for each V, D, J gene segment. In B cells, there are about 48 V, 20 D and six J functional gene segments. Identification of D gene segment in the V-D-J junction is rather difficult due to the short length of D gene segments and trimming at both ends of D gene segments during somatic recombination. Reads that were not mappable or that were mapped to non-functional gene segments were filtered out.

## Acknowledgements

We thank Dr. Aaron Logan and Dr. David B. Miklos for providing the immune repertoire sequencing data of patients as well as patient clinical data. We thank Dr. Carlos D. Bustamante and Dr. Hua Chen for helpful discussion of population genetic modeling. This work was supported by NIH P01-HG000205 to Ronald W. Davis, and NIH grant GM28016 to Marcus W. Feldman.

## Supporting Information

**S1 Fig.**
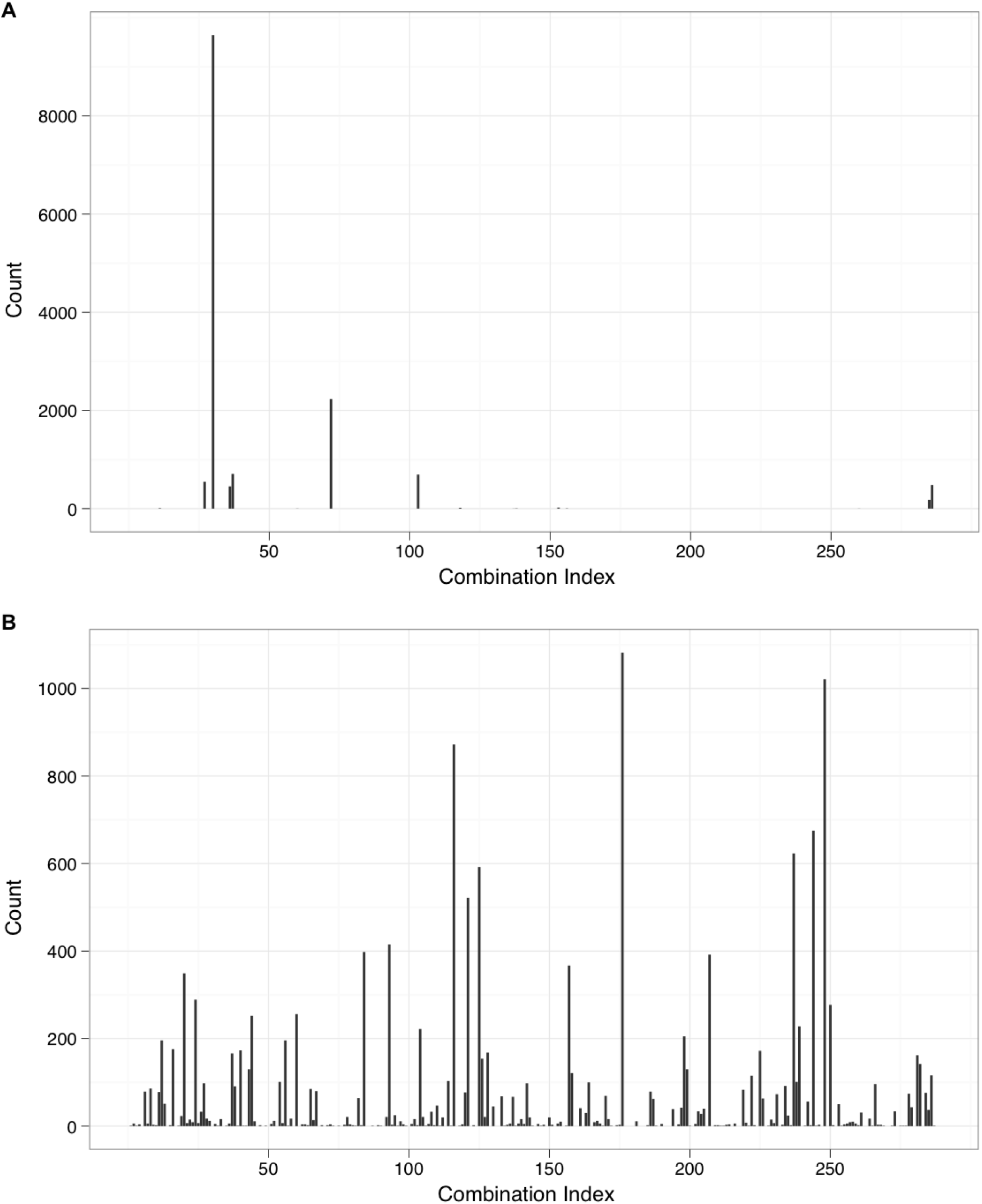
Frequency spectra of V-J combinations by forward simulation with incorporation rates *θ* = 1 (A) and *θ* = 40 (B) across 288 V-J combinations assuming a population size of 15,000.

